# Functional diversification of the MADS-box gene family in fine-tuning the dimorphic transition of *Talaromyces marneffei*

**DOI:** 10.1101/2024.12.16.628698

**Authors:** Xueyan Hu, Yun Zhang, Juan Wang, Minghao Du, Yang Yang, James J. Cai, Ence Yang

**Affiliations:** Department of Microbiology & Infectious Disease Center, School of Basic Medical Sciences, Peking University Health Science Center, Beijing 100191, China; Department of Medical Bioinformatics, School of Basic Medical Sciences, Peking University Health Science Center, Beijing 100191, China; Department of Biochemistry and Molecular Biology, Peking University Health Science Center, Beijing 100191, China; Department of Veterinary Integrative Biosciences, Texas A&M University, College Station, Texas, United States of America

**Keywords:** *Talaromyces marneffei*, MADS-box, dimorphism transition, RNA-seq, ChIP-seq, fungal adaptation

## Abstract

The dynamic transition between yeast and hypha is a crucial adaptive mechanism for many human pathogenic fungi, including *Talaromyces marneffei*, a thermodimorphic fungus responsible for causing fatal talaromycosis. In the current study, we elucidated the roles of the MADS-box gene family in fine-tuning the dimorphic transition in *T. marneffei* through functional diversification. Utilizing adaptive laboratory evolution, we identified an enrichment of MADS-box genes in mutants deficient in yeast-to-mycelium transition. Further phylogenetic analyses revealed a significant expansion of MADS-box gene family within *T. marneffei.* Functional genetic manipulations revealed that overexpression of *mads9*, as opposed to its paralog *mads10*, effectively impeded the hyphal-to-yeast transition. Through integrating RNA sequencing (RNA-seq) and chromatin immunoprecipitation sequencing (ChIP-seq), we demonstrated that *mads9* and the previously characterized *madsA* modulated the rate of hyphal-to-yeast conversion by orchestrating metabolic pathways and membrane dynamics, respectively, with mutual regulation via shared target genes. Our findings illuminated the distinct functional roles of the MADS-box family in regulating dimorphic transitions in *T. marneffei*, offering new insights into fungal adaptability.

## Introduction

Human pathogenic fungi, predominantly environmental in origin, often undergo morphological transitions between yeast/yeast-like and hypha/hyphal-like forms to adapt to the human body temperature during infection processes^1,2^. Understanding the genetic mechanisms behind these transitions cross environments and hosts can provide valuable insights into managing current fungal pathogens and preventing the emergence of new ones^3,4^. *Talaromyces marneffei* (previously known as *Penicillium marneffei*), is an opportunistic human pathogen that poses a significant health risk in immunocompromised individuals, particularly in Southeast Asia and South China^5^. Unlike the other species in its genus, *T. marneffei* is capable of thermal dimorphism, switching between saprophytic hyphae and pathogenic yeasts forms in response to temperature changes^6^. Since these morphological shifts can be fully induced by temperature in the laboratory^7^, *T. marneffei* serves as an ideal model for studying the mechanisms of heat adaptation in human pathogenic fungi.

Previous studies have demonstrated that a crucial aspect of thermal adaptation in *T. marneffei* is its temperature-regulated dimorphic transition between hyphal and yeast forms^1^. This complex phenotypic change involves alterations in multiple genes and pathways, highlighting the intricate regulatory mechanisms at play^6^. Among the key regulators identified are the *abaA*^8^, *areA*^9^, *hgrA*^10^, *madsA*^11^ and *msgA*^12^ genes. During the hypha-to-yeast transition, mutations in the ATTS transcription factor gene *abaA* can lead to growth defects when conidia produced by hyphae convert to yeast. Conversely, during the yeast-to-hypha transition, overexpression of the C2H2 transcription factor gene *hgrA* inhibits yeast growth at 37°C, while overexpression of the MADS-box transcription factor gene *madsA* promotes this conversion at 37°C. During yeast morphogenesis *in vivo*, the GATA factor gene *areA* and the Dbl homology/BAR domain gene *msgA* facilitate or maintain yeast cell growth within the host. Notably, among these genes, *madsA* is the member of the MADS-box family, which has been extensively studied for its regulatory roles in plants. However, the role of the MADS-box family in fungal morphogenesis remains underexplored.

The MADS-box gene family consists of transcription factors that are prevalent in eukaryotic organisms^13^. This family is named after its initial members: MCM1 from *Saccharomyces cerevisiae*^14^, AGAMOUS from *Arabidopsis thaliana*^15^, DEFICIENS from *Antirrhinum majus*^16^, and SRF from *Homo sapiens*^17^. All MADS-box members share a conserved domain that is crucial for nuclear localization, DNA binding, protein dimerization, and interactions with accessory factors^18–20^. MADS-box transcription factors participate in a spectrum of biological processes. In plants and animals, these factors regulate key cell proliferation and differentiation, and are significant for muscle morphogenesis^21–24^. Notably, a large number of MADS-box transcription factors in flowering plants interact with each other and constitute a complex regulatory network that determine floral organ morphology^25^. In fungi, such as *Schizosaccharomyces pombe*, *Aspergillus fumigatus* and *Candida albicans*, several MADS-box genes are involved in cellular morphogenesis, responses to environmental signals, and maintenance of cell wall integrity^26–29^. Our previous studies on *T. marneffei* identified and validated the function of the *madsA* gene^11,30^,. Additionally, we observed significant gene duplication within the MADS-box transcription factor family, highlighting the need for further investigation into how this family contributes to the dimorphic transition.

In this study, through temperature-induced adaptive laboratory evolution we significantly enriched the MADS-box family genes in strains with impaired hyphal formation. We then generated overexpression and gene knockout mutants of these MADS-box genes using genetic manipulation, confirming that their distinct regulatory roles in the biphasic conversion of *T. marneffei*. Through RNA sequencing (RNA-seq) and chromatin immunoprecipitation sequencing (ChIP-seq), we found that the MADS- box transcription factors *mads9* and *madsA* regulate different downstream processes via distinct TF binding sites, while also sharing common target genes in fine-tuning dimorphic transition. The functional divergence of *mads9* and *madsA* emphasize the importance of the MADS-box family in understanding morphological transformation and adaptive evolution of *T. marneffei*.

## Results

### The MADS-Box gene family is Enriched in Dimorphic Transition-Deficient Mutants Derived from Adaptive Laboratory Evolution in *Talaromyces marneffei*

To better understand the regulatory mechanisms underlying dimorphic transition, we first generated a population of transition-deficient mutants by employing adaptive laboratory evolution (ALE) with morphotype separation (**Figure 1A**-**B**). Utilizing bulked segregant analysis (BSA) combined with high-throughput sequencing, we identified 426 single-nucleotide variants (SNVs) and 200 indels from the transition- deficient population (**Supplemental Figure 1A**). The distribution of mutation frequency indicated that the transition-deficient population was composed of multiple mutants rather than a single mutated strain (**Supplemental Figure 1B**). Among the 31 completely mutated sites, there were only six mutants resulted in alterations to the protein coding sequences, including three frameshift and three nonsynonymous variants, with half being related to transmembrane transport (**Supplemental Table 1**).

**Figure 1.**
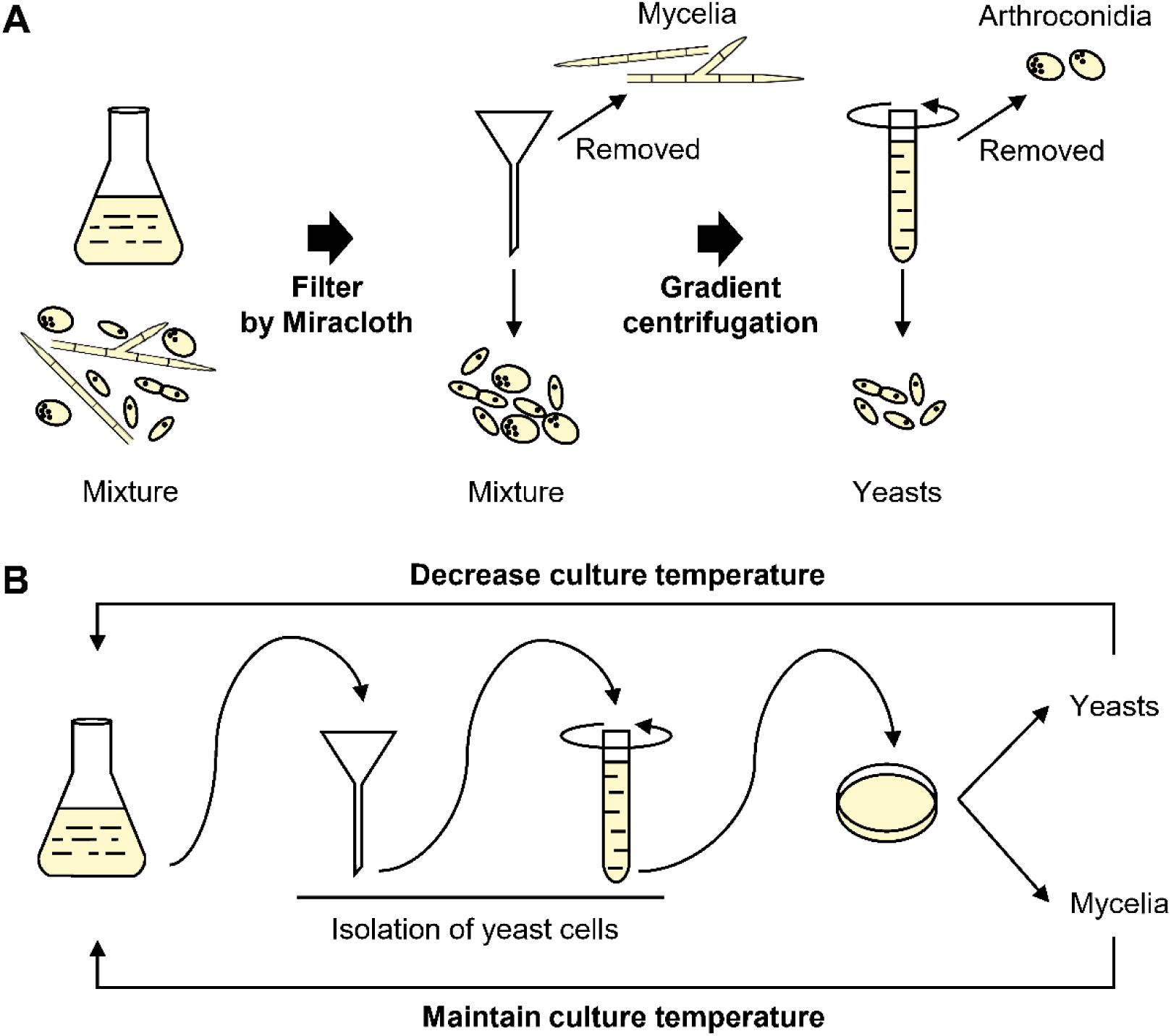
| Adaptive Laboratory Evolution Inducing Dimorphic Transition Defective Strains of *T. marneffei* PM1. A) Schematic workflow for isolating three different morphological cell types: mycelium, yeast, and conidia. Mycelium was filtered out using four layers of Miracloth, followed by centrifugation to separate yeast cells from conidia. B) Flowchart of the experimental evolution inducing dimorphic transition defective strains of *T. marneffei*. Cells from the SDB liquid medium were collected, and yeast cells were isolated using the method depicted in Figure A before being inoculated onto SDA plates to confirm their growth morphology. If substantial mycelium was present, cultures were maintained at the original temperature in SDB; if the predominant form was yeast, the incubation temperature was lowered by 1°C, continuing in SDB. This selection process was repeated until the culture temperature was reduced to 25°C.

We identified 792 structural variations (SVs), with five exhibiting mutation frequencies exceeding 80% (**Supplemental Figure 1C**). Given that the dimorphism- defective phenotype suggests a loss of function, we focused on the impact of three deletions among these high-frequency variants and identified 54 deleted genes. Functional enrichment analysis of the deleted genes revealed significant associations with GO functions such as protein dimerization activity (*P* = 4.0×10^-4^), protein kinase activity (*P* = 1.8×10^-3^) and DNA binding (*P* = 3.1×10^-3^). Interestingly, the MADS-box transcription factor superfamily (*P* = 1.6×10^-5^) was significantly enriched in IPR domain, with MADS genes (*mads9*, *mads10*, and *mads13*) identified within the deletion fragments. Combined with our previous findings^11,30^, we hypothesize that multiple members of the MADS-box family may be involved in the dimorphic transition of *T. marneffei*.

### Phylogenetic Analysis of the MADS-box Gene Family in *Talaromyces* Genus

To explore the potential regulatory role of the three identified MADS-box genes above (*mads9*, *mads10*, and *mads13*) in dimorphic transition, we performed a comprehensive phylogenetic analysis of the MADS-box gene family across seven *Talaromyces* species (**Supplemental Table 2**). Through the IPR protein domain database, we identified a total of 38 genes containing at least one MADS-box domain across genomes of these species. The *T. marneffei* PM1 strain exhibited the highest number with 15 MADS-box family genes, followed by *Talaromyces stipitatus* with 7. *Talaromyces islandicus*, *Talaromyces verruculosus*, and *Talaromyces atroroseus* each possessed 4 genes, while *Talaromyces cellulolyticus* and *Talaromyces amestolkiae* each contained 2 genes (**Figure 2A and Supplemental Table 2**).

**Figure 2.**
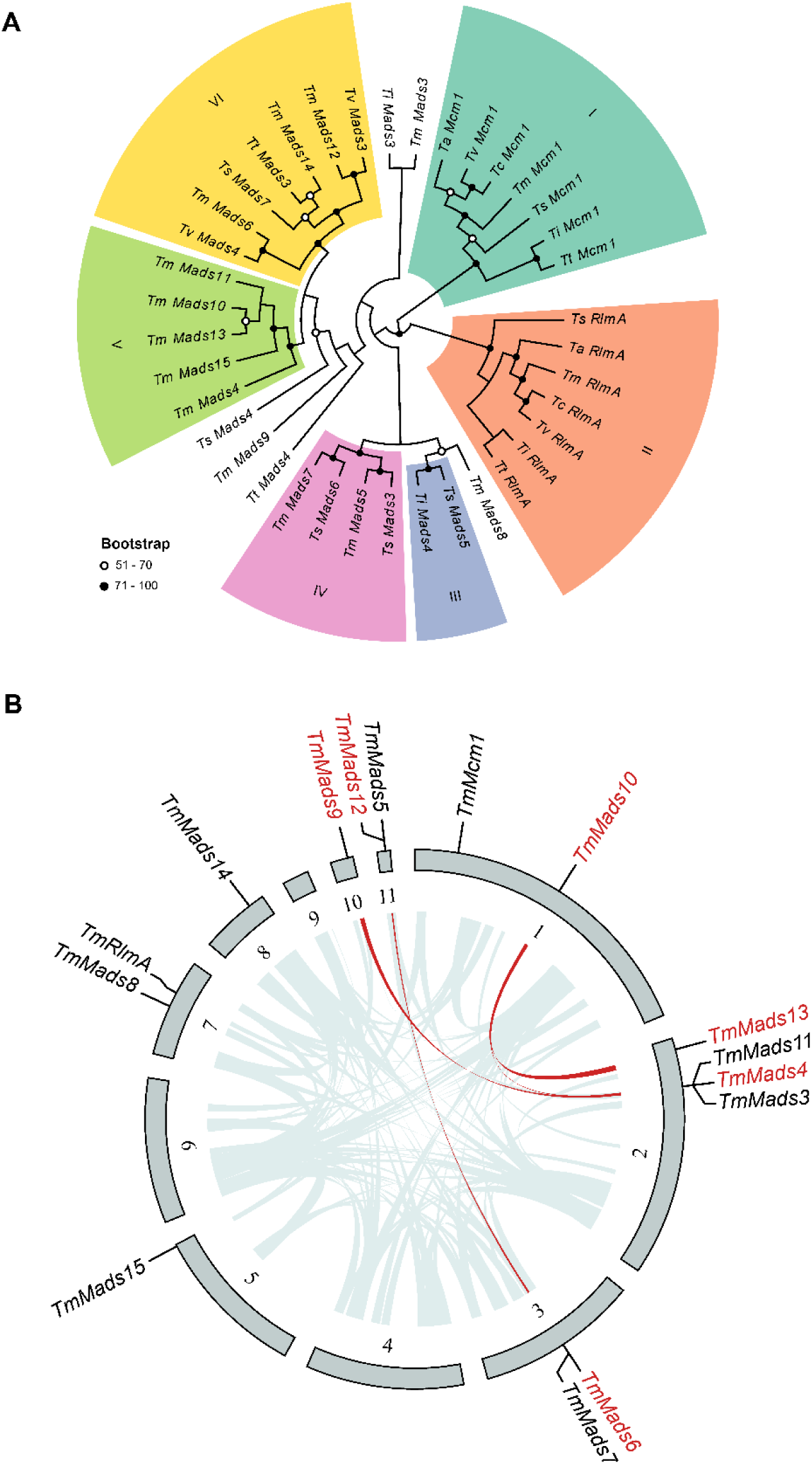
| Phylogenetic Analysis and Synteny of the MADS-box Family in *T. marneffei* A) Phylogenetic tree of the MADS-box gene family in the *Talaromyces* genus. B) Synteny analysis of the MADS-box gene family in *T. marneffei*. The outer layer represents the genomic sequences of wild-type strain PM1, with the locations of all MADS-box transcription factors annotated. The inner layer indicates the syntenic regions of the PM1 genome, with red highlighting the syntenic regions containing MADS-box transcription factors.

Phylogenetic analysis grouped 33 MADS-box genes into six main distinct clades (**Figure 2A**). Clades I and II corresponded to the highly conserved eukaryotic MADS-box transcription factors *Mcm1*^14^ and *RlmA*^28^. In the *T. marneffei* PM1 strain, five genes—*mads4*, *mads10*, *mads11*, *mads13*, and *mads15*—formed a monophyletic clade, suggesting a latest common ancestry. Further analysis of the *T. marneffei* PM1 strain revealed several collinear regions, including those between the *mads4* and *mads10* (eight gene pairs), *mads10* and *mads13* (six gene pairs), *mads4* and *mads9* (seven gene pairs), and *mads6* and *mads12* (six gene pairs) (**Figure 2B**).

Based on these collinear regions, the mechanisms responsible for the expansion of the MADS-box gene family in *T. marneffei* were inferred by MCScanX-transposed^31^. The species-specific increase in MADS-box transcription factors appeared to occur through two primary mechanisms: segmental duplication and transposition-mediated duplication. For instance, m*ads4*, *mads9*, *mads10*, and *mads13* may have originated from the segmental duplication of a common ancestral gene, while *mads6* and *mads12* likely arose from the segmental duplication of another ancestral gene. In contrast, the three genes *mads11*, *mads14*, and *mads15* seem to have resulted from independent transposition-mediated duplication events. Although *mads9* did not form a monophyletic branch with *mads10* and *mads13*, it is hypothesized that these three genes may have originated from the segmental duplication of a shared ancestral sequence (**Figure 2B**). This suggests that segmental duplication has played a pivotal role in the diversification and expansion of the MADS-box gene family, contributing to the adaptive evolution of dimorphic transition in *T. marneffei*.

### MADS-box Family Regulate Morphogenesis of *T. marneffei* with Functional Differentiation

Based on the potential monophyletic origin of *mads9*, *mads10*, and *mads13*, we further investigated the roles of *mads9* and *mads10* in regulating the morphogenesis of *T. marneffei*, considering the low expression level of *mads13* in our previous data^32^. We first constructed overexpression mutants for *mads9* and *mads10* and analyzed their growth at 37°C and 25°C conditions, respectively (**Supplemental Figure 2A**). After four days after incubation (dai) at 37°C, colonies of the OE-*mads9* strain appeared fluffier and smoother, whereas those of the wild-type PM1 strain were generally flat with more spikes. By 7 dai, the OE-*mads9* strains exhibited more wrinkled colony surfaces than the wild-type PM1 (**Figure 3**). The above results suggest that high expression of *mads9* may induce abnormalities in yeast cell morphogenesis. However, OE-*mads10* strains showed no significant morphological changes in comparison with the wild-type strains.

**Figure 3.**
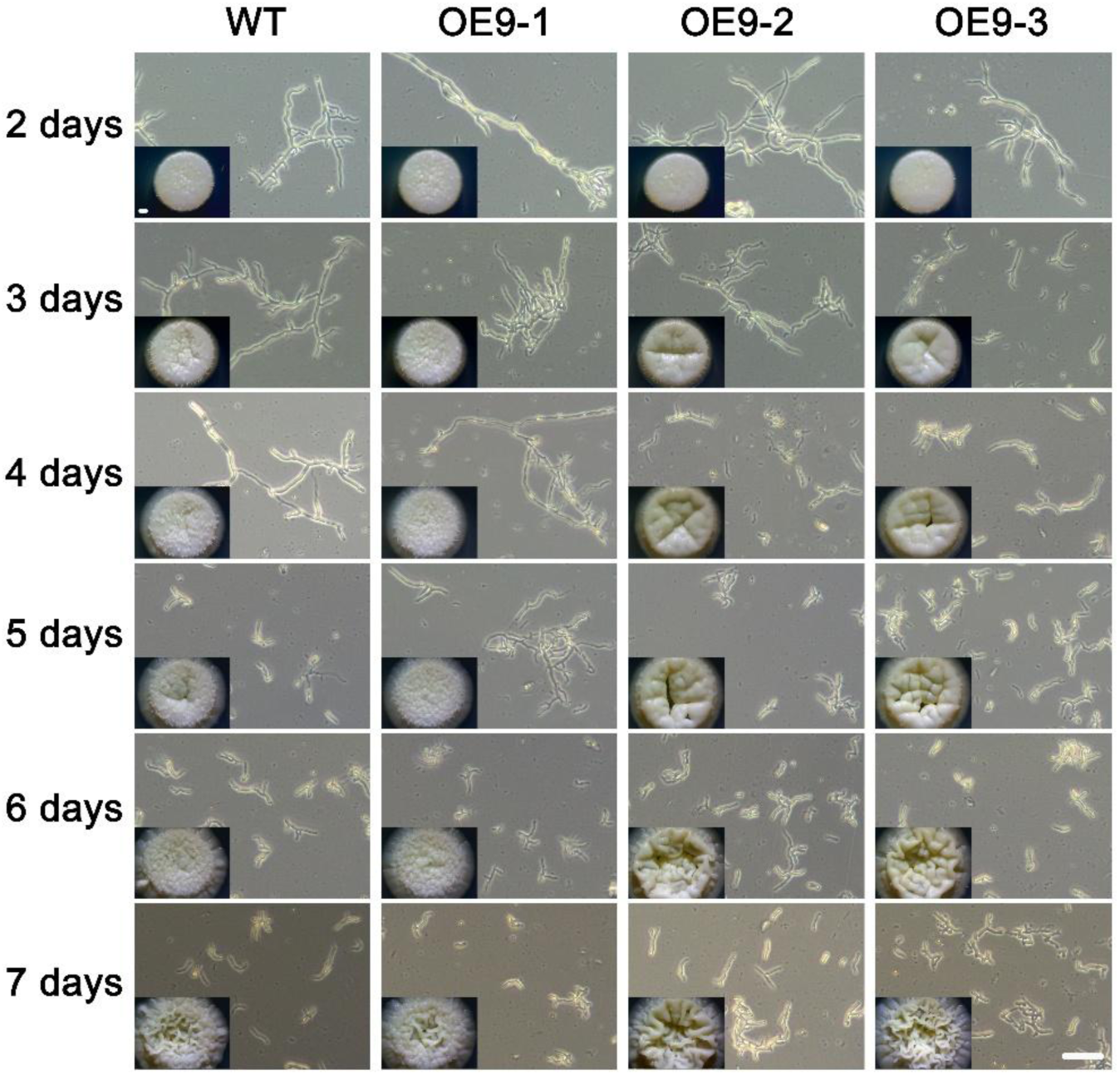
| Overexpression of *mads9* Promoted Morphogenesis of yeast cells in *T. marneffei*. Microscopic images were acquired under 20× objective lens with phase contrast effect. The morphogenesis process of yeast cells in OE-*mads9* strains seemed faster than in the wild-type PM1 strain. Bar = 20 μm. Time course phenotypes of colonies grown on SDA plates under 37°C were shown as insets. Bar = 1 mm.

Since the MADS-box gene family extensively regulates morphological developmental processes in plants^21^, we then explored whether the origin of this abnormal colony morphology was related to the morphological development process of yeast. Time course microscopic analysis revealed that the overall growth progression at 37°C of the OE-*mads9* strains were much faster than that of the wild-type PM1 cells.

By 4 dai, most of the wild-type PM1 cells exhibited elongated and branched hyphal-like growth, whereas abundant shorter and yeast-like cells were observed in the OE- *mads9* strains. In contrast, at 25°C, all strains grew in a filamentous form with normal spore formation as indicated by the yellow green colonies, and no obvious differences were observed between the wild-type and OE-*mads9* strains (**Supplemental Figure 3A**), indicating that the high expression level of *mads9* specifically interfered with the morphogenesis of yeast cells.

Finally, we generated knockout strains of *mads9* and *mads10* to explore the effects of loss of function of these genes (**Supplemental Figure 2B-C**). We monitored the growth of each genotype from 2 to 6 days post-inoculation. No significant phenotypic changes were observed in the KO-*mads9*, KO-*mads10*, and KO-*mads9/mads10* knockout lines at either 37°C or 25°C compared to the wild-type PM1 strain (**Supplemental Figure 3B-C**). This phenomenon may be due to the functional redundancy among MADS-box family genes^33^.

Together, these findings suggest that *mads9* could be influential in promoting yeast cell formation during constant growth at 37°C, while *mads10* has a much-limited effect.

### *Mads9* Regulate Dynamic Dimorphic Transition of *T. marneffei*

To further evaluate the role of MADS-box genes in dimorphic transitions, we conducted a mycelium-to-yeast (M-to-Y) transition experiment. However, at the colony level, there were no noticeable differences between the wild-type PM1 strains and the *mads9* knockout and overexpression strains (**Supplemental Figure 4A**). We then performed detailed microscopic analyses of the liquid mycelial cultures, focusing on changes at the cellular level. Initially, we examined the time-course morphological transformation of the wild-type PM1 strain under a microscope (**Figure 4A**). Before the M-to-Y transition began, most elongated mycelia exhibited typical even cell tips. At 2 hours post-transfer (hpt), many cell tips swelled like partially inflated balloons. By 3 hpt, these swollen regions expanded, with obvious branching at the mycelial ends and new tips forming around 6 hpt. Over time, these swollen regions continued to expand, becoming distinct from older, thinner mycelia. From 9 hpt onwards, swollen mycelial ends became predominant in the culture, with older reddish dying cells or debris easily distinguished. At 24 hpt and beyond, differential interference contrast (DIC) microscopy revealed bumpy and irregular cell surfaces. Active vesicles and septation were visible within cells, with contents appearing condensed.

**Figure 4.**
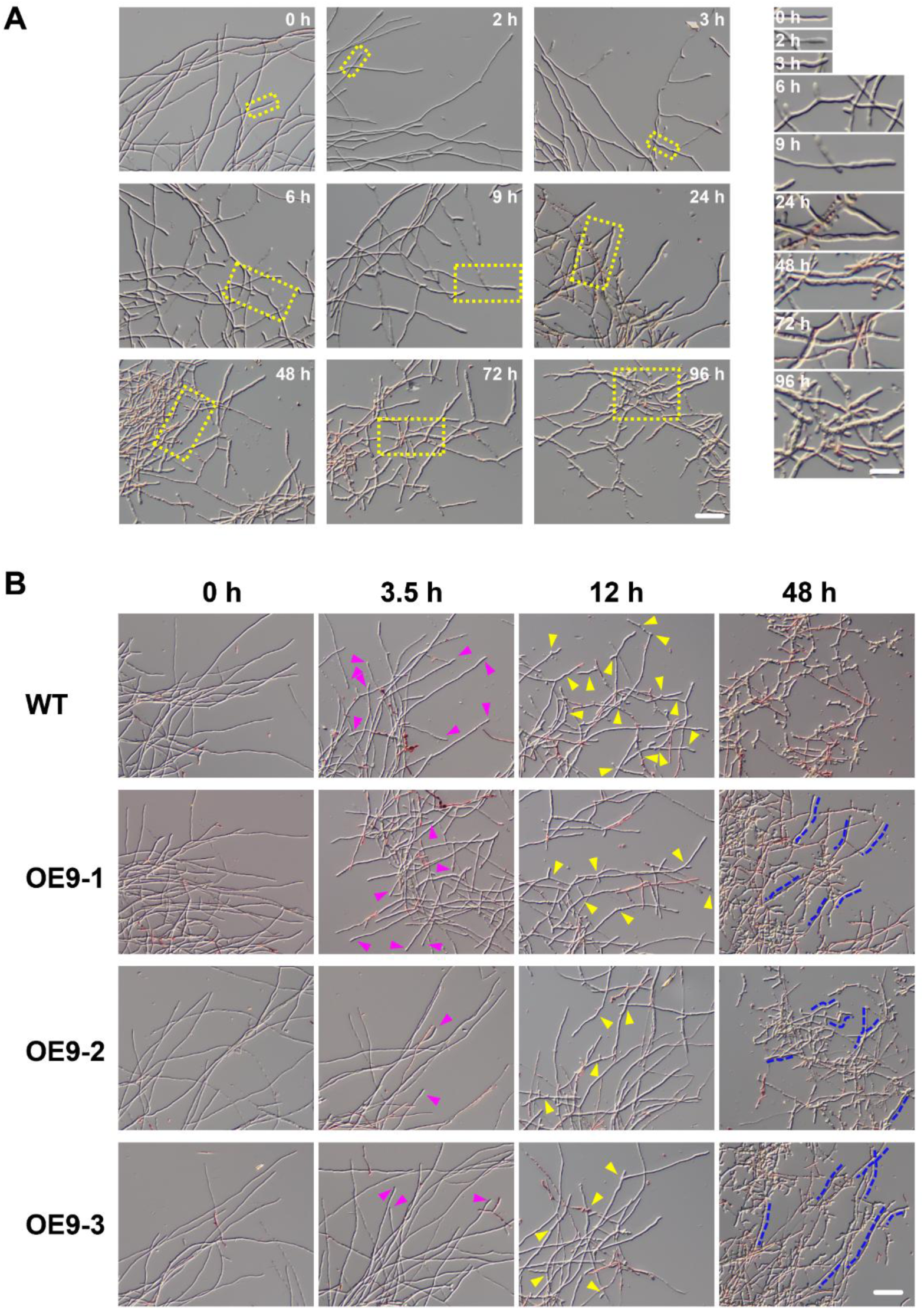
| Overexpression of *mads9* Delayed the M-to-Y Transition in *T. marneffei*. A) Time course microscopic analysis of the morphological changes in the wild-type cells grown in SDB transferred from 25°C to 37°C. Bar in the left panel is 50 μm. The highlighted region of interest in the left panel were magnified artificially and shown in the right panel. Bar = 25 μm. B) Microscopic analysis of the morphological changes in the wild-type and the OE-*mads9* cells grown in SDB transferred from 25°C to 37°C at indicated time points. Purple arrowheads indicated the swelling cells tips. Note that the ratio of swelling tips to the total cell ends invisible in the wild-type was bigger than those in the OE-*mads9* strains. Yellow arrowheads pointed towards the bumpy cell surfaces with newly-formed protrusions. Blue dotted lines labelled the longer cell remained in the OE-*mads9* strains. Bar = 50 μm.

Upon transition from 25°C to 37°C, OE-*mads9* strains showed a clear delay in the M-to-Y transition under the microscope (**Figure 4B**) though not in the colonies (**Supplemental Figure 4A**). At 3.5 hpt, fewer cells exhibited swelling tips in OE-*mads9* strains compared to wild-type. By around 12 hpt, most wild-type cell ends displayed active branching with numerous new tips forming; however, OE-*mads9* strain cell tips only began significant swelling at this point. As the transition progressed to around 48 hpt, condensed cells with bumpy surfaces interspersed with dying cells were observed in wild-type strains. In contrast, OE-*mads9* cells remained elongated and less condensed, indicating a delay compared to wild-type PM1 cells. Although the colonies of the KO-*mads9* strain showed no obvious phenotype (**Supplemental Figure 4A**), there was a certain degree of lag in the late stage (48 hpt) of the dynamic conversion process (**Supplemental Figure 4B**).

Overall, these results indicate that *mads9* was involved in regulating the dimorphism of *T. marneffei*, where high expression induces aberrant morphogenesis and delayed dimorphic transition.

### The MADS-box Gene Family Fine-Tunes Dimorphic Transitions Through Functional Differentiation

Given that both *mads9* and our previously studied *madsA*^11^ regulate dimorphic transitions with opposing trends, we next investigated the regulatory mechanisms of the MADS-box gene family to enhance our understanding of their roles in heat adaptation of *T. marneffei*. We performed RNA sequencing on the *mads9* and *madsA* mutants, which exhibited the most significant phenotypes, as well as on the wild-type PM1 strain under both mycelial and yeast growth conditions. Hierarchical clustering revealed consistent gene expression patterns across biological replicates, indicating high reproducibility of the sequencing data (**Figure 5A**).

**Figure 5.**
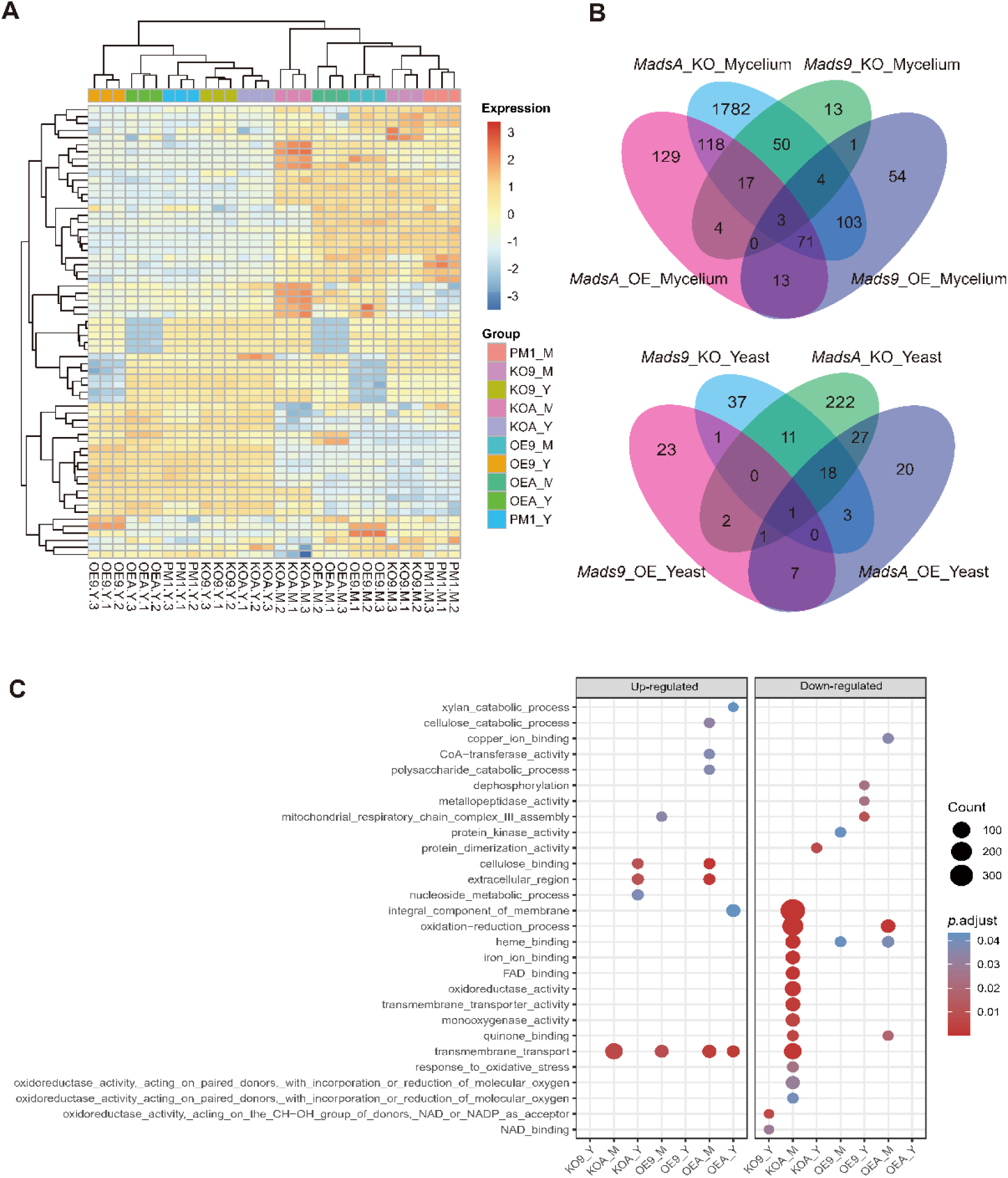
| Differential Gene Expression Analysis of MADS-box Mutant Strains in *T. marneffei* A) RNA-seq gene expression clustering of mutant and wild-type strains. The color code of the heatmap represents the *z*-scored gene expression level. The correlation of gene expression patterns and levels between biologically repeated samples was consistent. M, mycelium; Y, yeast. B) Differential gene expression analysis of *madsA* and *mads9* mutant strains. The Venn diagram displayed the count of unique and common differentially expressed genes (DEGs) between mutant strains and PM1 under two conditions. C) Enrichment results of differentially expressed genes in the mycelial and yeast phase. The size of point represents the count of DEGs, and the gradual color change from red to blue represents the adjust *P* value change from low to high.

Under hyphal conditions, differential expression analysis using DESeq2 identified 249, 92, 355, and 2148 differentially expressed genes (DEGs) in the OE-*mads9*, KO- *mads9*, OE-*madsA*, and KO-*madsA* strains, respectively **(Figure 5B)**. Gene Ontology (GO) enrichment analysis provided insights into the biological processes associated with the differentially expressed genes. In the *mads9* mutants, enriched GO terms included "transmembrane transport", "protein kinase activity" and "mitochondrial respiratory chain complex III assembly". Meanwhile, the *madsA* mutants were enriched for terms related to "membrane integral components", "oxidation-reduction processes" and "iron ion binding" (**Figure 5C**). Under yeast conditions, 35, 71, 77, and 282 DEGs were identified in the OE-*mads9*, KO-*mads9*, OE-*madsA*, and KO-*madsA* strains, respectively (**Figure 5B**). In this condition, the *mads9* mutants were enriched for terms such as "mitochondrial respiratory chain complex III assembly", "dephosphorylation", "NAD binding" and "metallopeptidase activity". On the other hand, the *madsA* mutants showed enrichment in terms including "transmembrane transport", "protein dimerization activity" and "cellulose binding" (**Figure 5C**).

These results indicated that the overexpression of the *mads9* gene may impact energy metabolism and cellular signaling pathways, thereby inhibiting the transition from hyphal to yeast form. On the other hand, *madsA* overexpression promotes hyphal development by modulating membrane integrity, redox balance, and ion homeostasis. This gene family expansion allows for the fine-tuning of the dimorphic transition in *T. marneffei*, adapting the fungus to different growth forms.

### Regulation Of Common Dimorphic Transition Effector Genes Between MADS-Box Families

Since *mads9* and *madsA* exhibit opposing regulatory effects by regulating distinct pathways, we further investigated whether there is potential crosstalk by performing ChIP-seq on FLAG-tagged *mads9* and *madsA* mutants under both hyphal and yeast conditions. The majority of *madsA* binding peaks were in promoter regions (82% in hyphal and 74% in yeast), while *mads9* showed significant binding in both promoter (60% in hyphal and 56% in yeast phases) and exon regions (**Figure 6A**). Relative to the transcription start site, the binding peaks of *mads9* and *madsA* were mainly distributed within 1 kb around the transcription start site (**Figure 6B**). Motif analysis identified enriched binding motifs consistent with genome-wide transcription factor binding profiles in both growth conditions (**Table 1**).

**Figure 6.**
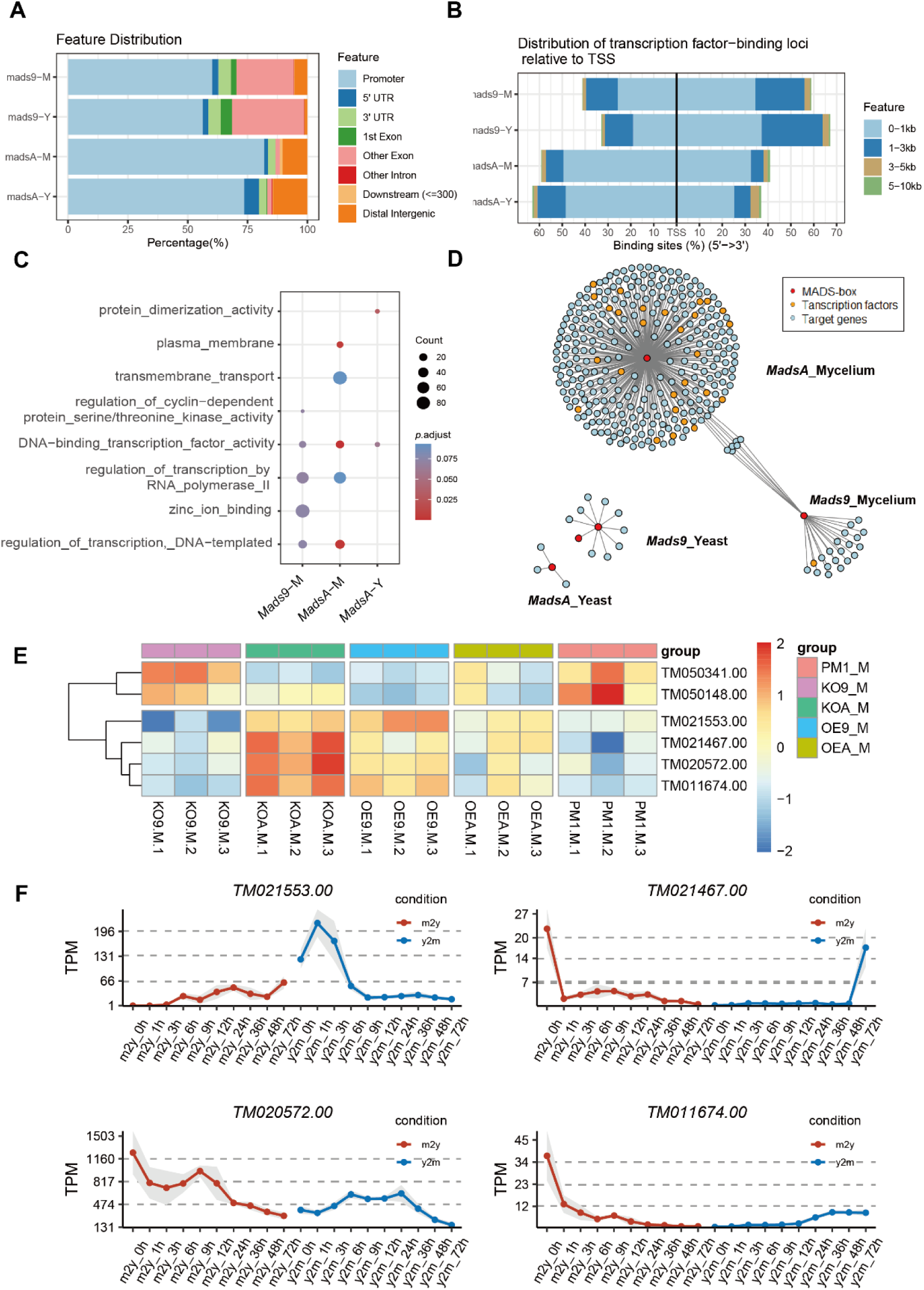
**| Gene Regulatory Network of the MADS-box Family.** A) Genome-wide binding patterns of MADS-box transcription factors. The ChIP-seq peaks are divided into several categories: promoter, first exon, other exons, first intron, other introns, 5’ untranslated region (5’ UTR), 3’ untranslated region (3’ UTR), downstream intergenic (within 300 kb of the 3’ UTR), and distal intergenic (>300 kb of the 3’UTR) regions. B) Distribution of MADS-box transcription factor binding sites relative to transcription start sites. C) Functional enrichment of target genes of MADS-box transcription factors. The size of point represents the count of target genes, and the gradual color change from red to blue represents the adjust *P* value change from low to high. D) Downstream regulatory network of MADS-box transcription factors. Each point represents a gene, with colors indicating the group to which the gene belongs: MADS-box transcription factors, other transcription factors, or additional genes. The lines illustrate the regulatory relationships between gene pairs. E) Heatmap of six genes were found to be co-regulated by *mads9* and *madsA*, with the *z*-score normalized expression levels indicated by the color bar (red indicating a relative higher expression level and blue indicating a lower expression level). F) Expression levels of four be co-regulated genes during dynamic dimorphic transition. Red represents the transformation from the mycelial phase to the yeast phase (M to Y), and blue represents the transformation from the yeast phase to the hyphal phase (Y to M). The dots represent the normalized mean expression of three biological replicates at the same time, and the shading represents the standard error range. The horizontal axis is time points. TPM, transcripts per million.

**Table 1.**
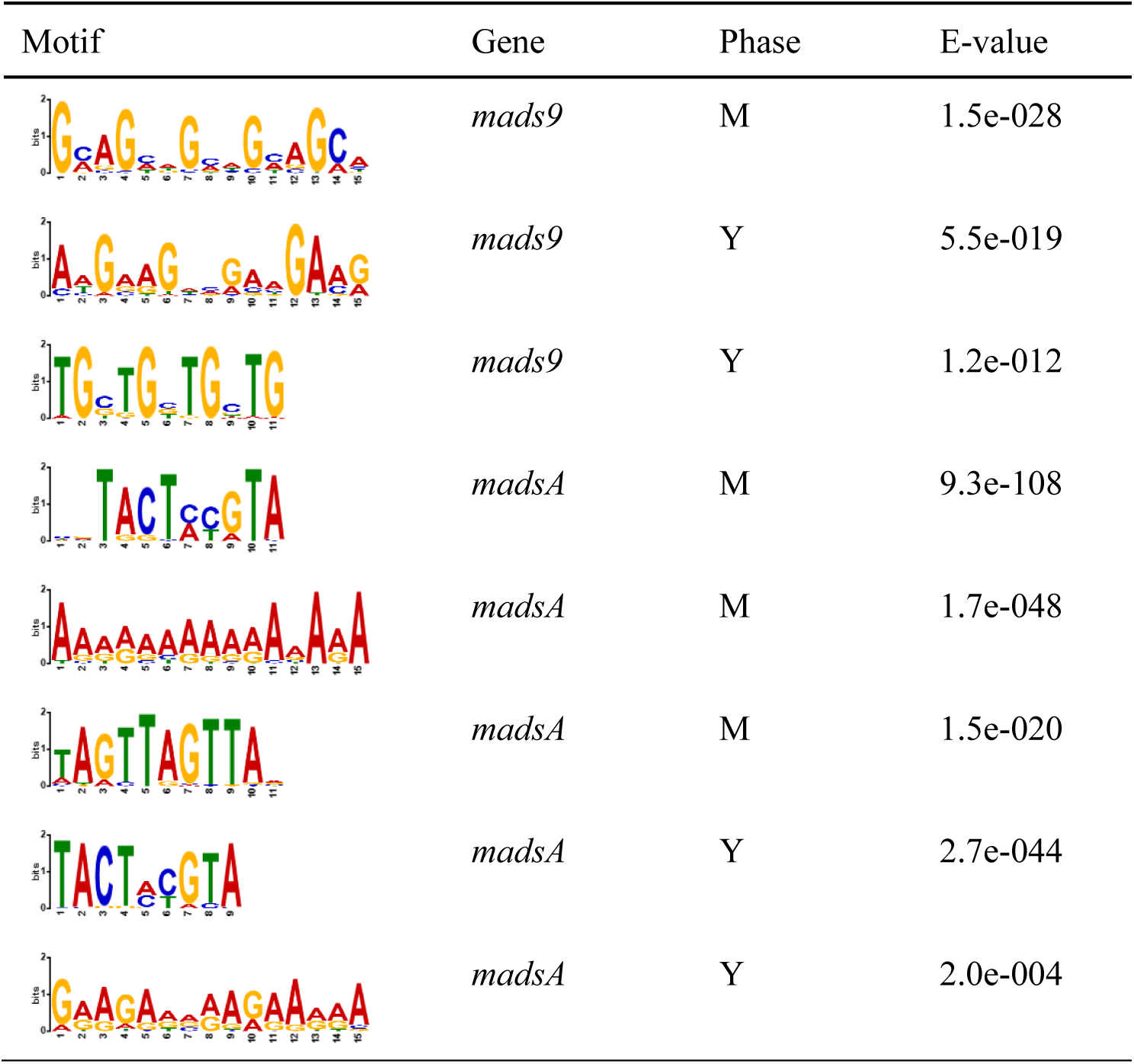
Enrichment motifs for binding sites of MADS-box transcription factors.

| Motif | Gene | Phase | E-value |
| --- | --- | --- | --- |
|  | *mads9* | M | 1.5e-028 |
|  | *mads9* | Y | 5.5e-019 |
|  | *mads9* | Y | 1.2e-012 |
|  | *madsA* | M | 9.3e-108 |
|  | *madsA* | M | 1.7e-048 |
|  | *madsA* | M | 1.5e-020 |
|  | *madsA* | Y | 2.7e-044 |
|  | *madsA* | Y | 2.0e-004 |

In the hyphal conditions, *mads9* target genes were enriched in "DNA binding transcription factor activity (GO:0003700)" and "zinc ion binding (GO:0008270)," while *madsA* targets were enriched in "DNA binding transcription factor activity (GO:0003700)" and "plasma membrane (GO:0005886)" (**Figure 6C**). However, in the yeast conditions, significant enrichment was not observed for *mads9* targets, whereas *madsA* targets were enriched in "protein dimerization activity (GO:0046983) "(**Figure 6C**), aligning with above RNA-seq results.

Integrating the ChIP-seq and RNA-seq data revealed a set of downstream genes regulated by *madsA* and *mads9*. Specifically, under hyphal conditions, 296 genes were regulated by *madsA* and 26 by *mads9*, while in the yeast phase, 3 genes were regulated by *madsA* and 11 by *mads9* (**Figure 6D**). Additionally, *Mp1p*-like protein 13 (*TM090004.00*) was identified as a downstream target of *mads9* in yeast. Given that *Mp1p* is a known virulence factor in *T. marneffei*^34^, this suggests that *mads9* may play a role in the pathogenesis of the fungus.

We then constructed the regulatory networks associated with *madsA* and *mads9* in both hyphal and yeast forms to infer pathway crosstalk (**Figure 6D**). We noticed six genes were found to be co-regulated by *mads9* and *madsA* under hyphal conditions (**Figure 6E**), including pathogenesis-related protein 5 (*TM021467.00*), aspergillopepsin-2 precursor (*TM021553.00*), monosaccharide transporter (*TM050148.00*), general amino acid permease (*TM050341.00*), and two hypothetical proteins (*TM020572.00* and *TM011674.00*). Among them, four of the co-regulated genes showed the same trend in OE-*madsA* and KO-*mads9*, and they showed significant differential expression in the time-series transition data (**Figure 6F**), suggesting that these genes may be downstream targets of *mads9* and *madsA* in regulating dimorphic transitions in different directions.

## Discussion

In this work, we demonstrated that the MADS-box gene family plays a pivotal role in the morphological transitions of *T. marneffei*, an opportunistic human pathogen capable of thermal dimorphism. Through laboratory evolution combined with high-throughput sequencing, phylogenetic analysis and genetic manipulation, and ChIP-seq, we identify *mads9* and *madsA* as key regulators in the temperature-induced dimorphic transition and show that they are functionally oppositely divergent.

### MADS-Box Family Regulation of Dimorphic Transition of *T. marneffei*

The ability of *T. marneffei* to switch from saprophytic hyphal forms to pathogenic yeast forms in response to temperature fluctuations is a sophisticated adaptation mechanism^6^. This study identifies the MADS-box transcription factors could fine-tuning this morphological transition by distinct genes. Specifically, the overexpression of *mads9* could block the mycelium-to-yeast transition processes at elevated temperatures, while the overexpression of *madsA* could promote the same transitions. In contrast, *mads10* had limited functions in dimorphic transition. This aligns with findings from other studies showing that MADS-box genes in fungi, such as *Candida albicans*, are implicated in morphogenesis and environmental response^26^. Furthermore, the interaction of these MADS-box genes with downstream processes, including transmembrane transport and redox reactions, underscores their multifaceted roles in fungal physiology and adaptability.

### Functional Divergence and Gene Evolution of MADS-Box Genes

The functional differences observed among the *mads9*, *mads10*, and *madsA* genes highlight the evolutionary dynamics within the MADS-box gene family. *Mads9* and *mads10* likely originated from a common ancestral gene, as evidenced by their structural similarities and the significant gene duplication observed within the family. However, their distinct regulatory functions in dimorphism reflect the process of neofunctionalization, where duplicated genes acquire new roles, thereby contributing to the organism’s adaptive potential.

In plants, similar patterns of gene duplication and functional divergence have been documented, particularly in the regulation of floral organ development^35–37^. This suggests that the evolutionary pressures acting on MADS-box genes in fungi may parallel those in plants, driving the diversification of regulatory functions in response to environmental challenges. The complex interplay among these genes could further elucidate the evolutionary trajectory of MADS-box transcription factors and their roles in fungal pathogenicity.

### Temperature Adaptation and Environmental-to-Pathogenic Fungal Transition

Thermodimorphism, where fungi switch between saprophytic hyphal growth at lower temperatures and yeast forms at human body temperature, is a critical adaptation that enables survival within the host^1^. The ability of *T. marneffei* to undergo a temperature-induced dimorphic transition is a hallmark of its pathogenic potential, mirroring a broader evolutionary strategy employed by environmental fungi as they switch to opportunistic human pathogens. In this study, we revealed that the MADS-box transcription factors, particularly *mads9* and *madsA* are involving in this transition with opposite functions. The enrichment of MADS-box genes in *T. marneffei* compared to other species within the *Talaromyces* genus suggests that gene duplication and functional diversification are important drivers of its ability to thrive in natural/host dual environments. This expands our understanding of how environmental fungi evolve to exploit host niches, a process that is becoming increasingly relevant as climate change alters ecosystems and increases human exposure to emerging fungal pathogens.

## Methods

### Strains and culture conditions

The wild-type *Talaromyces marneffei* strain PM1, isolated from a patient with culture-documented talaromycosis in Hong Kong^38^, was utilized in this study. All strains (both mutants and wild type) were cultured on Sabouraud Dextrose Agar (SDA, BD Difco^TM^) plates at 25°C or 37°C for indicated days, either to facilitate conidia collection or to phenotypic observation. For total RNA extraction, all strains were grown in Sabouraud Dextrose Broth (SDB, BD Difco^TM^) at 25°C or 37°C for three days before the next-step operation. For mycelium-to-yeast transition experiment, following 3 days of culture in SDB or 4 days on SDA plates at 25°C, samples were transferred to 37°C and continued growing. Time course observations were conducted at indicated time intervals after temperature transition. Microscopic analysis was performed on the Primo Star microscope (Zeiss) equipped with the 20× objective (Plan-ACHROMAT, NA0.4, Ph2) or the upright microscope (Olympus, BX53) equipped with the 20× objective (UPLFLN20X, NA0.5). Images were captured with ZEN software, and the cellSens Standard software bundled with Extend Focus Image (EFI) acquisition module (Olympus).

### Isolation of Mycelium, Yeast, and Conidia

The fungal suspension was filtered through four layers of Miracloth, and the filtrate was collected in a new 50 mL centrifuge tube. The suspension was centrifuged at 10,000 × g for 5 minutes at 4°C. The supernatant was carefully removed using a 10 mL disposable pipette, leaving approximately 1 mL of liquid at the bottom of the tube. The cells were resuspended by gently tapping the tube. A 15 mL centrifuge tube containing 13 mL of 1× PBS solution was prepared, and the cell suspension was slowly added along the wall of the tube to minimize disturbance. The mixture was centrifuged at 100 × g for 1 minute at 4°C, discarding the upper 12 mL of liquid and retaining 1 mL of concentrated yeast-form cells at the bottom.

### Adaptive Laboratory Evolution

The Talaromyces marneffei PM1 strain was inoculated into SDB liquid medium and incubated at 32°C with shaking at 180 rpm for 7 days. Yeast-form cells were collected and re-inoculated into fresh SDB medium under the same conditions for an additional 3 days, followed by further isolation. After each passage, 10 μL of the culture was plated onto SDA agar plates and incubated at the same temperature for 7 days to assess growth morphology. If the strain predominantly exhibited yeast-form growth, the incubation temperature was decreased by 1°C, repeating the aforementioned steps. When the temperature reached 25°C, a dimorphic transition defective strain was maintained in the yeast form at ambient temperature.

### Phylogenetic Analysis of MADS-box Transcription Factor Family

Based on IPR protein domain annotation results^39^, genes containing at least one MADS-box transcription factor-related domain (IPR002100, IPR033896, IPR033897, IPR036879) were screened from seven species of basket fungi. The identified MADS-box genes were subjected to multiple sequence alignment using MAFFT software^40^ (v7.407). Subsequently, a phylogenetic tree based on maximum likelihood was constructed using RAxML software^41^ (v8.2.12), with 1,000 bootstrap replicates and a Γ distribution for site rate variation, automatically selecting the amino acid substitution model based on the alignment results. The phylogenetic tree was visualized using EvolView software^42^ (v2).

### Genome Synteny Analysis

The protein sequences of T. marneffei were aligned with those of itself and six other basket fungi using BLASTp software^43^, applying an E-value threshold of < 1 × 10^-5^. Synteny regions in the PM1 strain’s genome were detected using MCScanX-transposed software^31^, which also inferred the types of gene duplication within the genome. A Python script was used to organize synteny regions and identify those containing MADS-box transcription factors. The results of the genome synteny analysis were visualized using Circos software^44^ (v0.69).

### Preparation of Gene Overexpression Strains

Genomic DNA was extracted from *T. marneffei*, and high-fidelity PCR was employed to amplify fragments of the coding region of the target gene for overexpression. The fragment of the promoter of the *gpdA* gene and the hygromycin resistance gene were amplified from the pEGFP-1 plasmid using high-fidelity PCR. These fragments were then assembled into an overexpression vector using the ClonExpress II One Step Cloning Kit (Vazyme). Strains were transformed with overexpression plasmids following a previously described protocol with modifications^45^. Briefly, strains were cultured on SDA medium at 25°C for 7 days. Spores were collected by scraping with a fine toothpick, washed twice, and incubated for germination at 37°C with shaking at 150 rpm for 40 hours. Germinated spores were centrifuged at 4,200 × g for 5 minutes, and excess supernatant was removed. Germinated spores were resuspended in 40 mL of enzymatic hydrolysis solution (β-Glucuronidase and Driselase, Sigma) at 37°C with shaking at 80 rpm until approximately 50% protoplasts were produced. The protoplasts were filtered using Miracloth (Calbiochem, Germany) and washed twice. Clean protoplasts were stored at -80°C until transformation. For transformation, a total of 5 μg of plasmid DNA was mixed with 1 × 10^6 protoplasts and incubated on ice for 30 minutes. Subsequently, 1 mL of PTC solution (60% PEG6000, 100 mM Ca^2+^) was added, and the mixture was incubated at room temperature for 60 minutes. Transformants were cultured on selective medium at 37°C for mutant selection and validated through whole genome sequencing.

### Preparation of Gene Knockout Strains

The *mads9* knockout strains were acquired mainly based on the published method^46^. In brief, the whole knockout constructs, including the Cas9 constructs, the gRNA constructs and the gene deletion constructs were electroporated into *T. marneffei* and the transformants were subsequently selected from colonies grown at 37°C. The Cas9 cassette was amplified from the vector pCas9P2AEGFP using primers FragC9-F and FragC9-R. The gene specific gRNA designed with the CRISPR gRNA (guide RNA) Design Tool for Eukaryotic Pathogens (http://grna.ctegd.uga.edu/) were ligated into the pTmCas9 vector generated in our previous study^47^, and the gRNA cassette was amplified using primer pairs TmU6P-F and Scaffold-R. The gene deletion constructs included the use of homologous arms (around 1.3 kb) with split marker gene confers resistance to drug G418. In the 1^st^ round of PCR, the 5’ homologous arm, the selection marker, and the 3’ homologous arm were amplified. In the 2^nd^ round of PCR, the 5’ and the 3’ arms were fused with the selection marker. In the 3^rd^ round of PCR, the full-length deletion construct was amplified. The *mads10* knockout constructs contained only the *mads10* deletion constructs including 5’ and 3’ homologous arms (around 1 kb) with split hygromycin (HYG) marker gene. For single gene knockout transformations, the wild-type PM1 strain was used. For *mads9* and *mads10* double gene knockout, the *mads10* knockout constructs were transformed into the KO-*mads9* strains. Primers used were indicated in the Supplemental Table 3. The knockout constructs were electroporated into competent *T. marneffei* conidia cells using Gene Pulser Xcell Total System (BIO-RAD). The positive transformants appeared on SDA plates supplemented with 200 μg/mL G418 or hygromycin at 37°C after growth for 2-3 days. For competent *T. marneffei* conidia cell preparation, the conidia cells pre-germinated in SDB for about 12-13 h at 25°C were collected initially. Then the cells underwent sequential wash with once of chilled ddH2O and twice of 1M sorbitol (Sigma-Aldrich^®^), and suspended with 1 M sorbitol at concentrations above 10^9^ cells/mL finally.

### Verification of Gene Expression

For relative gene expression analysis, the cDNA was synthesized from total RNA (1 μg) using HiScript III 1st Strand cDNA Synthesis Kit (Vazyme, China). Diluted cDNA templates were amplified in ChamQ Universal SYBR qPCR Master Mix (Vazyme, China) using specific primer pairs designed for specific genes as listed in Supplemental Table 3. The actin gene was used as an internal control. Three technical replicates in each test were carried out. Due to the difficulties for designing a primer pair which specifically targeted *mads10*, genotyping PCR using primer pair 10up-F and 10down- R could only be used for the *mads10* knockout strain identification. The positive *mads10* knockout transformant with selective marker inserted should produce a 4.3bp DNA fragment, whereas the wild-type produced a 2.9kb DNA fragment.

### RNA-seq Library and Data Analysis

Total RNA of strains was extracted using the E.Z.N.A. fungal RNA kit (Omega Bio- Tek), following the manufacturer’s instructions with DNase I digestion to eliminate genomic DNA. The RNA concentration was measured using Nanodrop (Thermo Fisher Scientific Inc. USA), and the quality was assessed using Qsep1 (BiOptic, Inc., New Taipei City, Taiwan). RNA with a RIN > 8 was selected and stored at -80℃ until downstream preparation. RNA-seq library was performed and sequenced on the Illumina XTEN platform by at the Novogene Bioinformatics Institute (Beijing, China), generating paired-end 150 bp reads. Adapters and low-quality segments were trimmed from the raw sequencing data. The cleaned data were aligned to the PM1 reference genome^32^ using HISAT2^48^, and gene expression TPM (Transcripts per million) matrices were generated with StringTie2^49^, correcting for sequencing depth and gene length.

Differential expressed genes (DEGs) were identified by DESeq2 package^50^ with FDR < 0.05 and |log2(FC)| > 1. Functional enrichment of genes were preformed using ClusterProfiler package^51^.

### ChIP-seq Library and Data Analysis

Strains were collected and frozen by immersion in liquid nitrogen. Frozen samples were ground and cross-linked with 1% formaldehyde for 10 min at room temperature with gentle rotation. Cross-linking was stopped by adding 200 mM final concentration of glycine solution in PBS and cells were rotated at room temperature for 5 min. Fixed cells were pelleted at 1500 ×*g* for 10 min at 4℃ and washed twice with 10 mL of ice-cold PBS and pelleted at 1500 ×*g* for 10 min at 4℃ each time. Washed cell pellet was then stored at -80℃ until further processing. ChIP-seq was performed using Anti-FLAG (DYKDDDDK Tag Rabbit mAb, CST#14793) at the Boyun Huakang Gene Technology Ltd (Beijing, China). The clean reads generated by ChIP-seq were evaluated using FastQC (https://github.com/s-andrews/FastQC) and aligned to the reference PM1 genome with Bowtie2^52^. Samtools^53^ was then used to sort and index BAM files and uniquely mapped reads were extracted using Sambamba^54^. MACS2^55^ was employed to call peaks for each sample, using input DNA as a control. ChIP-seq peaks were annotated, compared and visualized using ChIPseeker^56^ package, and motif analysis was conducted using MEME^57^.

## Data analysis and visualization

All data analyses were performed using R (v4.1.2). Visualizations were generated using ggplot2 (v3.3.6), unless specified otherwise.

## Statistical tests

All statistical tests were conducted in R (v4.1.2), with the relevant test methods and statistics indicated in the corresponding sections of the text.

## Data Availability

The pooled sequencing data for the dimorphic transition mutant strains of *T. marneffei* have been uploaded to NCBI under the project number PRJCA002719. The RNA-seq and ChIP-seq were uploaded to NCBI under the access number GSE279912 and GSE279913.

## Acknowledgements

This work was funded by National Natural Science Foundation of China (31970008 and 32170091, Ence Yang).

## Author contributions

Yang Yang, James J. Cai and Ence Yang designed and supervised the project. Minghao Du, Yun Zhang and Xueyan Hu performed bioinformatics analysis. Juan Wang and Yun Zhang performed fungal experiments and molecular experiments. Xueyan Hu wrote the manuscript along with Ence Yang. All authors approved the final version of the manuscript.

---

**Supplemental Figure 1.**
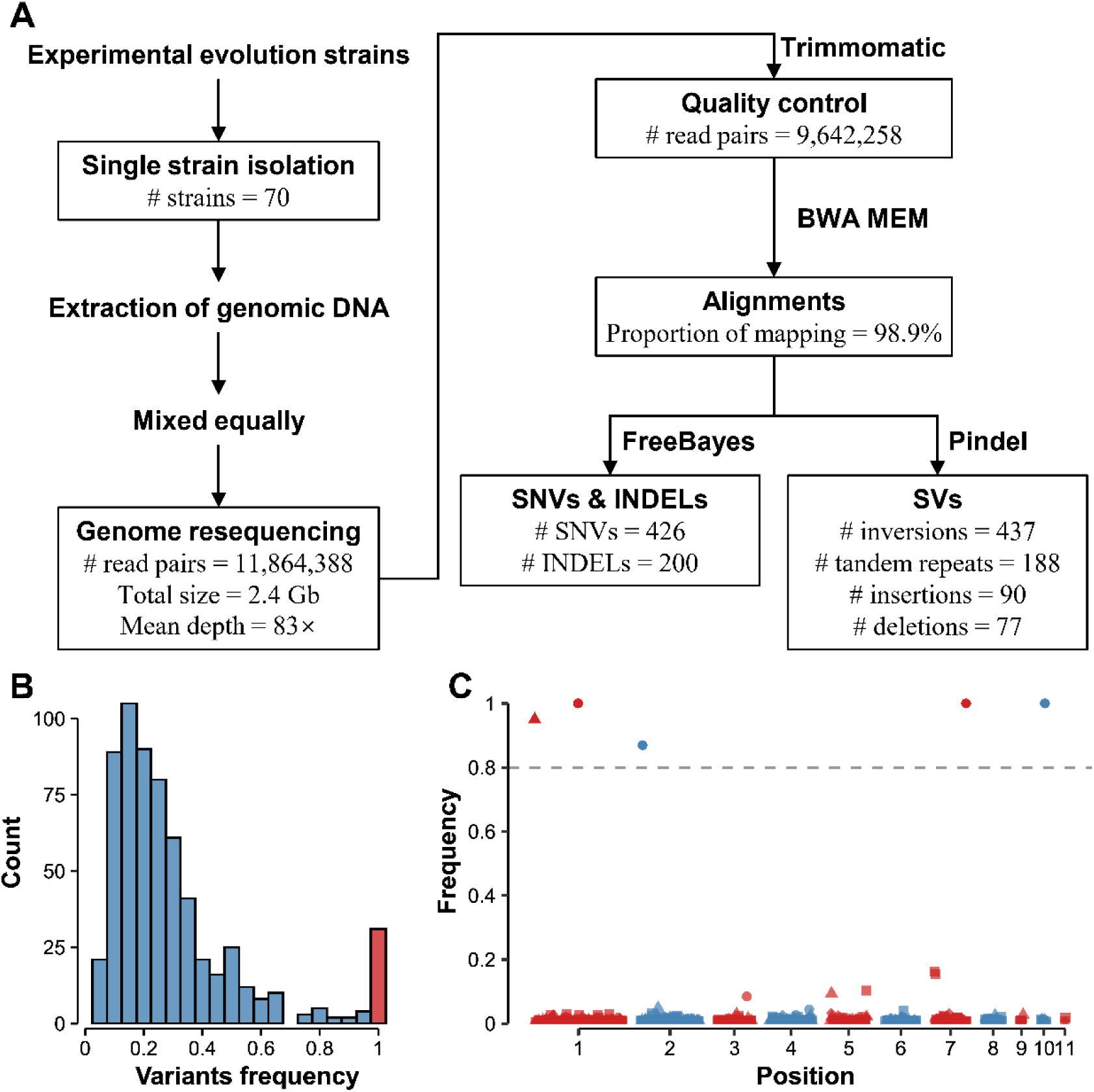
| Results of mixed group analysis of *T. marneffei* **dimorphism-defective population.** A) Flowchart of mixed group analysis of experimental evolution mutant population. B) Frequency distribution of SNVs and INDELs in the genome of experimental evolution mutant population. The horizontal axis represents the frequency of SNVs INDELs in the mutants, and the vertical axis represents the number of mutation sites contained in the corresponding frequency. The red bar represents the number of sites with a mutation frequency of 100%. C) Distribution of structural variation in the genome of experimental evolution mutants. The horizontal axis represents the position of the genome of *T. marneffei* PM1 strain, and adjacent genomic sequences are distinguished by red and blue. The vertical axis represents the frequency of structural variation in the mutants, and the gray dotted line represents the structural variation screening threshold (structural variation mutation frequency exceeds 80%). The shape of the scattered points represents the type of structural variation: rectangles represent inversion mutations, triangles represent tandem duplications, diamonds represent fragment insertions, and circles represent fragment deletions.

**Supplementary Figure 2.**
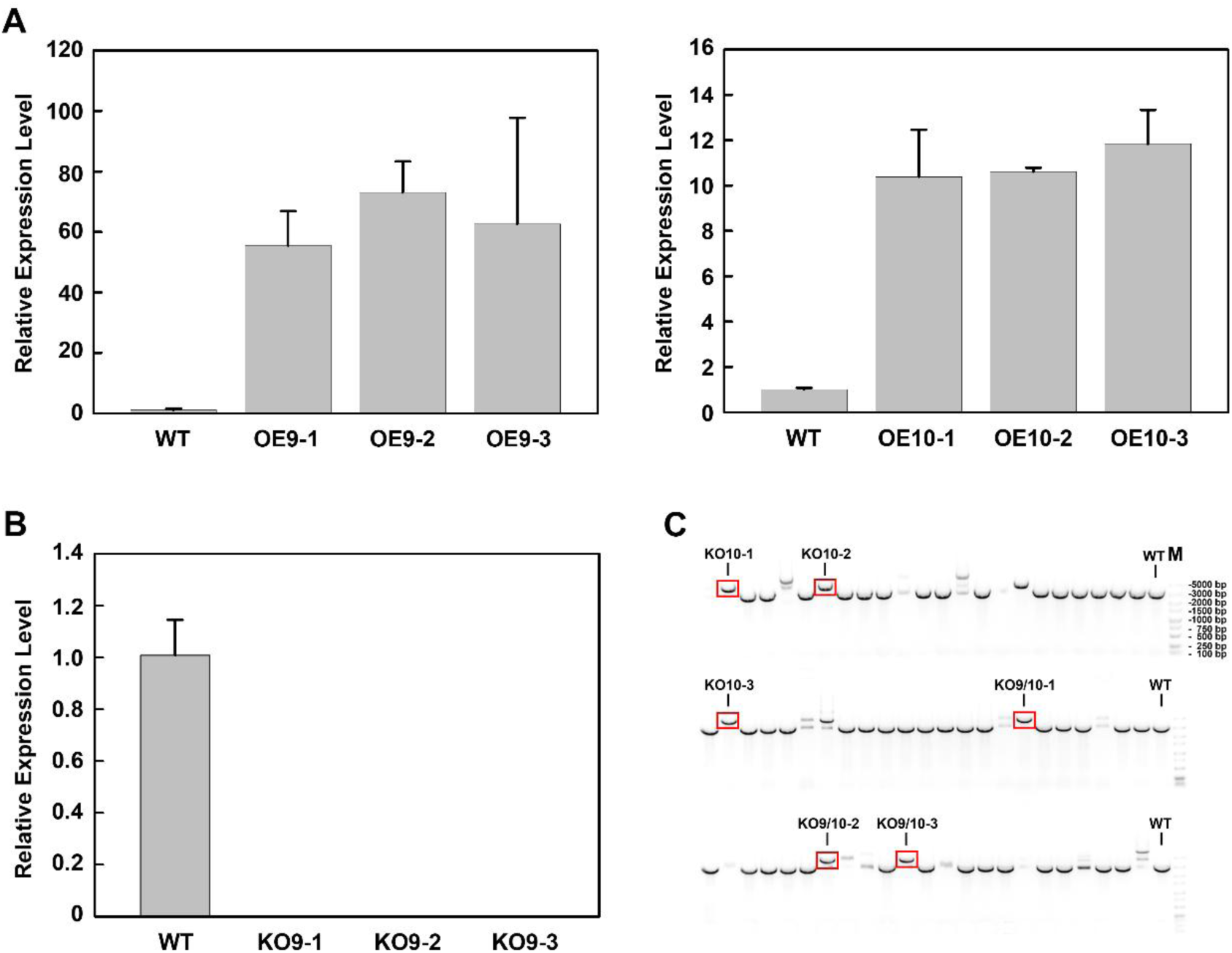
| Gene Expression Level Analysis of *T. marneffei* Strains. A) Relative expression level of *mads9* and *mads10* in the overexpression strains compared with the wild-type. B) Relative expression level of *mads9* in the knockout strains compared with the wild- type. Each of the test in qPCR included three technical replicates. Error bar represent mean±SE. C) Genotyping results revealed the positive transformants. The highlighted DNA band indicted the bigger DNA fragment amplified (around 4.3 kb), compared with the smaller DNA fragment amplified (around 2.9 kb) from the wild-type background.

**Supplemental Figure 3.**
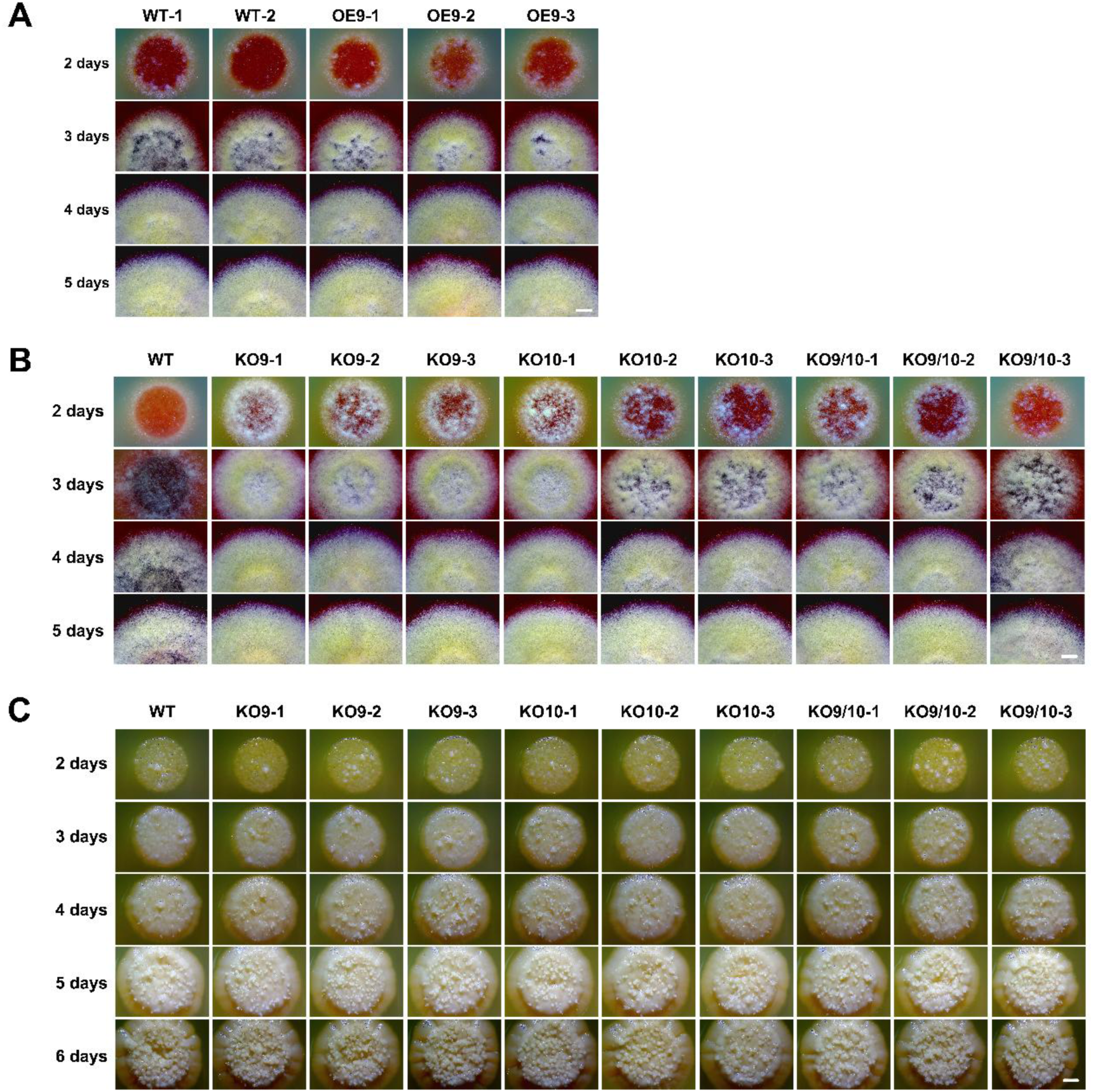
| The Phenotypes of *T. marneffei* Strains Grown on SDA Plates at Constant Temperature. A) Comparison of the OE-*mads9* and the wild-type colonies grown at 25°C. Bar = 2 mm. B) Comparison of the colonies of *mads9*, *mads10* and *mads9/mads10* knockout strains and the wild-type grown at 25°C. Bar = 2 mm. C) Comparison of the colonies of *mads9*, *mads10* and *mads9/mads10* knockout strains and the wild-type grown at 37°C. Bar = 2 mm.

**Supplemental Figure 4.**
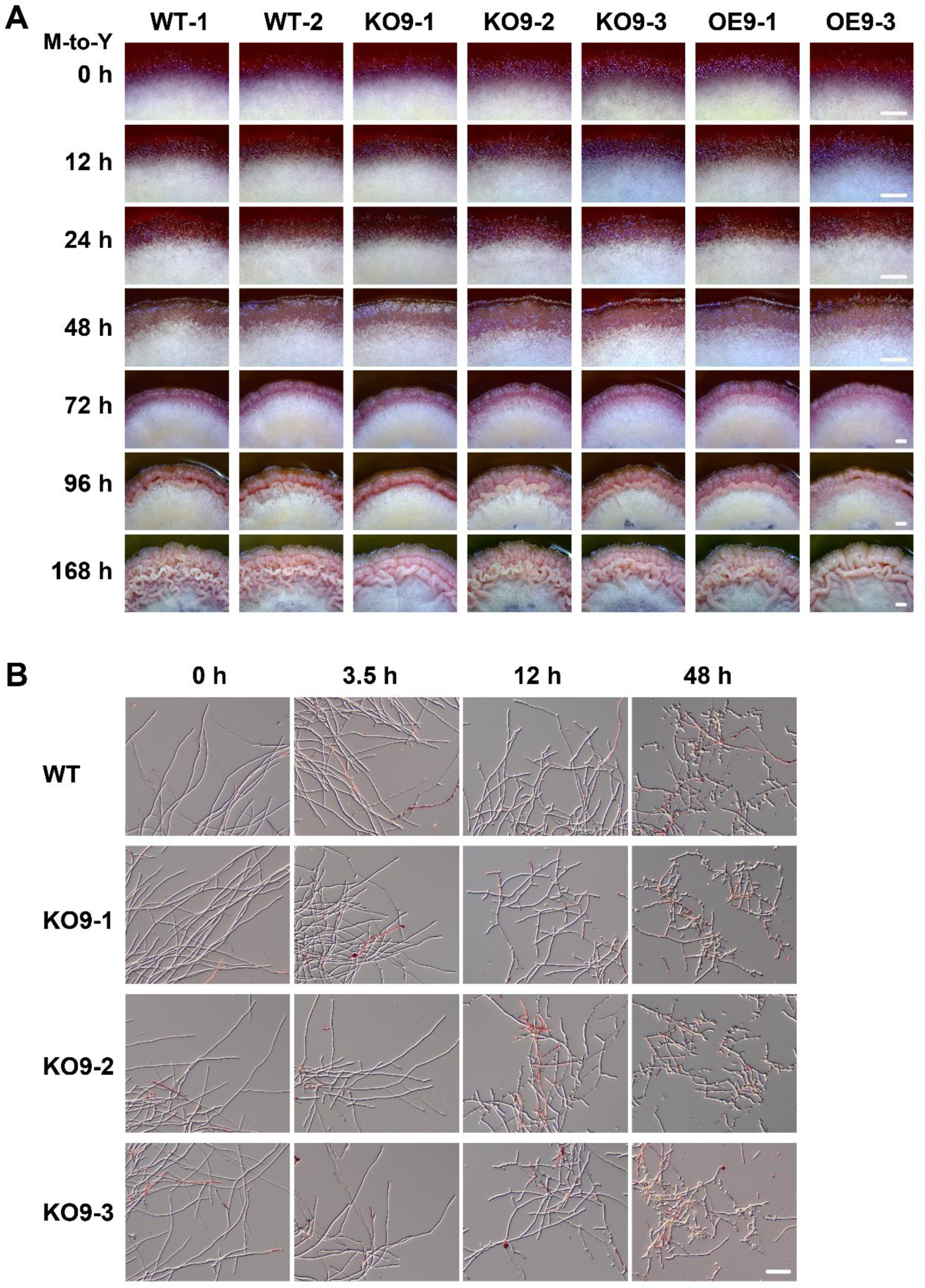
| The Morphological Changes of *T. marneffei* Strains During the M-to-Y Transition. A) Comparison of the colonies of *mads9* knockout and overexpression strains and the wild-type on SDA plates after transferred from 25°C to 37°C. Bar =1 mm. B) Microscopic analysis of the morphological changes in the wild-type and the KO-*mads9* cells grown in SDB transferred from 25°C to 37°C at indicated time points. Bar = 50 μm.

## Supplemental Tables

**Supplemental Table 1 |** Annotation of mutation sites in genes and their upstream and downstream regions for dimorphism-defective strains in adaptive laboratory evolution.

**Supplemental Table 2 |** Information on MADS-box family members in *Talaromyces* genus

**Supplemental Table 3 |** Primers used in this study.

## Reference

1. Boyce, K. J. & Andrianopoulos, A. Fungal dimorphism: the switch from hyphae to yeast is a specialized morphogenetic adaptation allowing colonization of a host. FEMS Microbiol. Rev. 39, 797–811 (2015).

2. Xiao, W. et al. Response and regulatory mechanisms of heat resistance in pathogenic fungi. Appl. Microbiol. Biotechnol. 106, 5415–5431 (2022).

3. Seidel, D. et al. Impact of climate change and natural disasters on fungal infections. Lancet Microbe 5, e594–e605 (2024).

4. Greenspan, S. E. et al. Constant-temperature predictions underestimate growth of a fungal amphibian pathogen under individual host thermal profiles. J. Therm. Biol. 111, 103394 (2023).

5. Nierman, W. C., Fedorova-Abrams, N. D. & Andrianopoulos, A. Genome sequence of the AIDS-sssociated pathogen *Penicillium marneffei* (ATCC18224) and its near taxonomic relative *Talaromyces stipitatus* (ATCC10500). Genome Announc. 3, e01559–14 (2015).

6. Wang, F., Han, R. & Chen, S. An overlooked and underrated endemic mycosis- Talaromycosis and the pathogenic fungus *Talaromyces marneffei*. Clin. Microbiol. Rev. 36, e0005122 (2023).

7. Andrianopoulos, A. Control of morphogenesis in the human fungal pathogen *Penicillium marneffei*. Int. J. Med. Microbiol. IJMM 292, 331–347 (2002).

8. Borneman, A. R., Hynes, M. J. & Andrianopoulos, A. The *abaA* homologue of *Penicillium marneffei* participates in two developmental programmes: conidiation and dimorphic growth. Mol. Microbiol. 38, 1034–1047 (2000).

9. Lau, S. K. P. et al. Proteome profiling of the dimorphic fungus *Penicillium marneffei* extracellular proteins and identification of glyceraldehyde-3-phosphate dehydrogenase as an important adhesion factor for conidial attachment. FEBS J. 280, 6613–6626 (2013).

10. Bugeja, H. E., Hynes, M. J. & Andrianopoulos, A. HgrA is necessary and sufficient to drive hyphal growth in the dimorphic pathogen *Penicillium marneffei*. Mol. Microbiol. 88, 998–1014 (2013).

11. Wang, Q. et al. MADS-Box transcription factor madsA regulates dimorphic transition, conidiation, and germination of *Talaromyces marneffei*. Front. Microbiol. 9, 1781 (2018).

12. Weerasinghe, H., Bugeja, H. E. & Andrianopoulos, A. The novel Dbl homology/BAR domain protein, MsgA, of *Talaromyces marneffei* regulates yeast morphogenesis during growth inside host cells. Sci. Rep. 11, 2334 (2021).

13. Ng, M. & Yanofsky, M. F. Function and evolution of the plant MADS-box gene family. Nat. Rev. Genet. 2, 186–195 (2001).

14. Passmore, S., Maine, G. T., Elble, R., Christ, C. & Tye, B. K. *Saccharomyces cerevisiae* protein involved in plasmid maintenance is necessary for mating of MAT alpha cells. J. Mol. Biol. 204, 593–606 (1988).

15. Yanofsky, M. F. et al. The protein encoded by the Arabidopsis homeotic gene agamous resembles transcription factors. Nature 346, 35–39 (1990).

16. Schwarz-Sommer, Z. et al. Characterization of the Antirrhinum floral homeotic MADS-box gene deficiens: evidence for DNA binding and autoregulation of its persistent expression throughout flower development. EMBO J. 11, 251–263 (1992).

17. Norman, C., Runswick, M., Pollock, R. & Treisman, R. Isolation and properties of cDNA clones encoding SRF, a transcription factor that binds to the c-fos serum response element. Cell 55, 989–1003 (1988).

18. Lamb, R. S. & Irish, V. F. Functional divergence within the APETALA3/PISTILLATA floral homeotic gene lineages. Proc. Natl. Acad. Sci. U. S. A. 100, 6558–6563 (2003).

19. Becker, A. & Theissen, G. The major clades of MADS-box genes and their role in the development and evolution of flowering plants. Mol. Phylogenet. Evol. 29, 464– 489 (2003).

20. Immink, R. G. H., Gadella, T. W. J., Ferrario, S., Busscher, M. & Angenent, G. C. Analysis of MADS box protein-protein interactions in living plant cells. Proc. Natl. Acad. Sci. U. S. A. 99, 2416–2421 (2002).

21. Castelán-Muñoz, N. et al. MADS-Box genes are key components of genetic regulatory networks involved in abiotic stress and plastic developmental responses in plants. Front. Plant Sci. 10, 853 (2019).

22. Marques, I. et al. Transcriptomic analyses reveal that *Coffea arabica* and *Coffea canephora* have more complex responses under combined heat and drought than under individual stressors. Int. J. Mol. Sci. 25, 7995 (2024).

23. Black, B. L. & Olson, E. N. Transcriptional control of muscle development by myocyte enhancer factor-2 (MEF2) proteins. Annu. Rev. Cell Dev. Biol. 14, 167–196 (1998).

24. Eisenmann, K. M. et al. 5q- myelodysplastic syndromes: chromosome 5q genes direct a tumor-suppression network sensing actin dynamics. Oncogene 28, 3429– 3441 (2009).

25. Herrera-Ubaldo, H. et al. The protein-protein interaction landscape of transcription factors during gynoecium development in Arabidopsis. Mol. Plant 16, 260–278 (2023).

26. Rottmann, M., Dieter, S., Brunner, H. & Rupp, S. A screen in *Saccharomyces cerevisiae* identified CaMCM1, an essential gene in *Candida albicans* crucial for morphogenesis. Mol. Microbiol. 47, 943–959 (2003).

27. Baker, C. R., Hanson-Smith, V. & Johnson, A. D. Following gene duplication, paralog interference constrains transcriptional circuit evolution. Science 342, 104–108 (2013).

28. Rocha, M. C., et al. *Aspergillus fumigatus* MADS-box transcription factor rlmA is required for regulation of the cell wall integrity and virulence. G3 Bethesda Md 6, 2983–3002 (2016).

29. He, Z. et al. Participation of a MADS-box transcription factor, Mb1, in regulation of the biocontrol potential in an insect fungal pathogen. *J. Invertebr.* Pathol. 170, 107335 (2020).

30. Yang, E. et al. Signature gene expression reveals novel clues to the molecular mechanisms of dimorphic transition in Penicillium marneffei. PLoS Genet. 10, e1004662 (2014).

31. Wang, Y., Li, J. & Paterson, A. H. MCScanX-transposed: detecting transposed gene duplications based on multiple colinearity scans. Bioinforma. Oxf. Engl. 29, 1458–1460 (2013).

32. Du, M. et al. Unraveling the dynamic transcriptomic changes during the dimorphic transition of *Talaromyces marneffei* through time-course analysis. Front. Microbiol. 15, 1369349 (2024).

33. De Bodt, S., Theissen, G. & Van de Peer, Y. Promoter analysis of MADS-box genes in eudicots through phylogenetic footprinting. Mol. Biol. Evol. 23, 1293–1303 (2006).

34. Narayanasamy, S. et al. A global call for talaromycosis to be recognised as a neglected tropical disease. Lancet Glob. Health 9, e1618–e1622 (2021).

35. Huu, C. N., Keller, B., Conti, E., Kappel, C. & Lenhard, M. Supergene evolution via stepwise duplications and neofunctionalization of a floral-organ identity gene. Proc. Natl. Acad. Sci. 117, 23148–23157 (2020).

36. Abraham-Juárez, M. J. et al. Evolutionary variation in MADS Box dimerization affects floral development and protein abundance in maize. Plant Cell 32, 3408– 3424 (2020).

37. Bowman, J. L. & Moyroud, E. Reflections on the ABC model of flower development. Plant Cell 36, 1334–1357 (2024).

38. Woo, P. C. Y. et al. Draft genome sequence of *Penicillium marneffei* strain PM1. Eukaryot. Cell 10, 1740–1741 (2011).

39. Apweiler, R. et al. The InterPro database, an integrated documentation resource for protein families, domains and functional sites. Nucleic Acids Res. 29, 37–40 (2001).

40. Katoh, K. & Standley, D. M. MAFFT multiple sequence alignment software version 7: improvements in performance and usability. Mol. Biol. Evol. 30, 772–780 (2013).

41. Stamatakis, A. RAxML version 8: a tool for phylogenetic analysis and post- analysis of large phylogenies. Bioinforma. Oxf. Engl. 30, 1312–1313 (2014).

42. Evolview v2: an online visualization and management tool for customized and annotated phylogenetic trees - PubMed. https://pubmed.ncbi.nlm.nih.gov/27131786/.

43. Camacho, C. et al. BLAST+: architecture and applications. *BMC Bioinformatics* **10**, 421 (2009).

44. Krzywinski, M. et al. Circos: an information aesthetic for comparative genomics. Genome Res. 19, 1639–1645 (2009).

45. Development of CRISPR-Cas9 genome editing system in Talaromyces marneffei. Microb. Pathog. 154, 104822 (2021).

46. Lin, J., Fan, Y. & Lin, X. Transformation of *Cryptococcus neoformans* by electroporation using a transient CRISPR-Cas9 expression (TRACE) system. Fungal Genet. Biol. FG B 138, 103364 (2020).

47. Zhang, X. et al. Development of CRISPR-Cas9 genome editing system in *Talaromyces marneffei*. Microb. Pathog. 154, 104822 (2021).

48. Kim, D., Langmead, B. & Salzberg, S. L. HISAT: a fast spliced aligner with low memory requirements. Nat. Methods 12, 357–360 (2015).

49. Kovaka, S. et al. Transcriptome assembly from long-read RNA-seq alignments with StringTie2. Genome Biol. 20, 278 (2019).

50. Love, M. I., Huber, W. & Anders, S. Moderated estimation of fold change and dispersion for RNA-seq data with DESeq2. Genome Biol. 15, 550 (2014).

51. Yu, G., Wang, L.-G., Han, Y. & He, Q.-Y. clusterProfiler: an R package for comparing biological themes among gene clusters. Omics J. Integr. Biol. 16, 284– 287 (2012).

52. Langmead, B. & Salzberg, S. L. Fast gapped-read alignment with Bowtie 2. Nat. Methods 9, 357–359 (2012).

53. Danecek, P. et al. Twelve years of SAMtools and BCFtools. GigaScience 10, giab008 (2021).

54. Tarasov, A., Vilella, A. J., Cuppen, E., Nijman, I. J. & Prins, P. Sambamba: fast processing of NGS alignment formats. Bioinforma. Oxf. Engl. 31, 2032–2034 (2015).

55. Feng, J., Liu, T., Qin, B., Zhang, Y. & Liu, X. S. Identifying ChIP-seq enrichment using MACS. Nat. Protoc. 7, 1728–1740 (2012).

56. Yu, G., Wang, L.-G. & He, Q.-Y. ChIPseeker: an R/Bioconductor package for ChIP peak annotation, comparison and visualization. Bioinforma. Oxf. Engl. 31, 2382– 2383 (2015).

57. Bailey, T. L., Johnson, J., Grant, C. E. & Noble, W. S. The MEME Suite. Nucleic Acids Res. 43, W39–49 (2015).

